# Where does the rainbow end? A case study of self-sustaining rainbow trout *Oncorhynchus mykiss* populations in tropical rivers

**DOI:** 10.1101/2022.06.21.497001

**Authors:** Marie Nevoux, Amandine D. Marie, Julien Raitif, Jean-Luc Baglinière, Olivier Lorvelec, Jean-Marc Roussel

## Abstract

The great physiological, behavioral and morphological plasticity of salmonids contributed to the success of their introduction into a wide range of habitats. However, rainbow trout (*Oncorhynchus mykiss*) is the only salmonid species to live in the tropics. Like for any cold-water species, water temperature is assumed to be a major constraint to the presence and expansion of rainbow trout. In this study, we collected detailed records of the introduction history of rainbow trout at 35 sites across the island over 80 years, and we document the presence of self-sustainable trout at each site. We explored the linear relationship between water temperature and elevation.

## INTRODUCTION

Native to the northern hemisphere, salmonid species have been introduced worldwide (Welcomme 1988; Crawford & Muir 2008; Lecomte *et al*. 2013), especially brown trout (*Salmo trutta*), brook trout (*Salvelinus fontinalis*), coho salmon (*Oncorhynchus kisutch*) and rainbow trout (*Oncorhynchus mykiss*). Their great physiological, behavioral and morphological plasticity contributes to the success of their introduction into a wide range of environmental conditions (Sauter *et al*. 2001). Since they are a cold-water species, however, successful introductions have been reported mainly at high latitudes and in temperate regions (Cowx 1997; Stanković *et al*. 2015; Koutsikos *et al*. 2019), since water temperature is a major constraint to their ontogeny and expansion. Among salmonids, rainbow trout has the widest latitudinal distribution in its native range (Fig. 1a), from 23-64°N (MacCrimmon 1971; Welcomme 1988; Crawford & Muir 2008). The population in the Sierra Madre Occidental Mountains of Mexico represents the southernmost native range of any salmonid (Behnke 2010), along with the Formosan landlocked salmon (*Oncorhynchus masou formosanus*) in Asia. These populations reflect rainbow trout’s ability to survive under extreme thermal conditions (MacCrimmon 1971; Gall & Crandell 1992). Its optimal range is 10-16°C, but adults can endure peaks of 25°C (Raleigh 1984; Jalabert & Fostier 2010) and even up to 28-30°C (Matthews & Berg 1997; Sloat & Osterback 2013). In contrast, it reproduces in winter to meet requirements for lower temperatures. Spawning and successful egg development have been recorded at temperatures of 4-13°C (Morrison & Smith 1986; Pankhurst *et al*. 1996; Jalabert & Fostier 2010) and even up to 15.5°C (Scott & Crossman 1973). Above 20°C, however, the fecundity rate and embryonic survival decrease greatly (King *et al*. 2003).

**Fig. 1.**
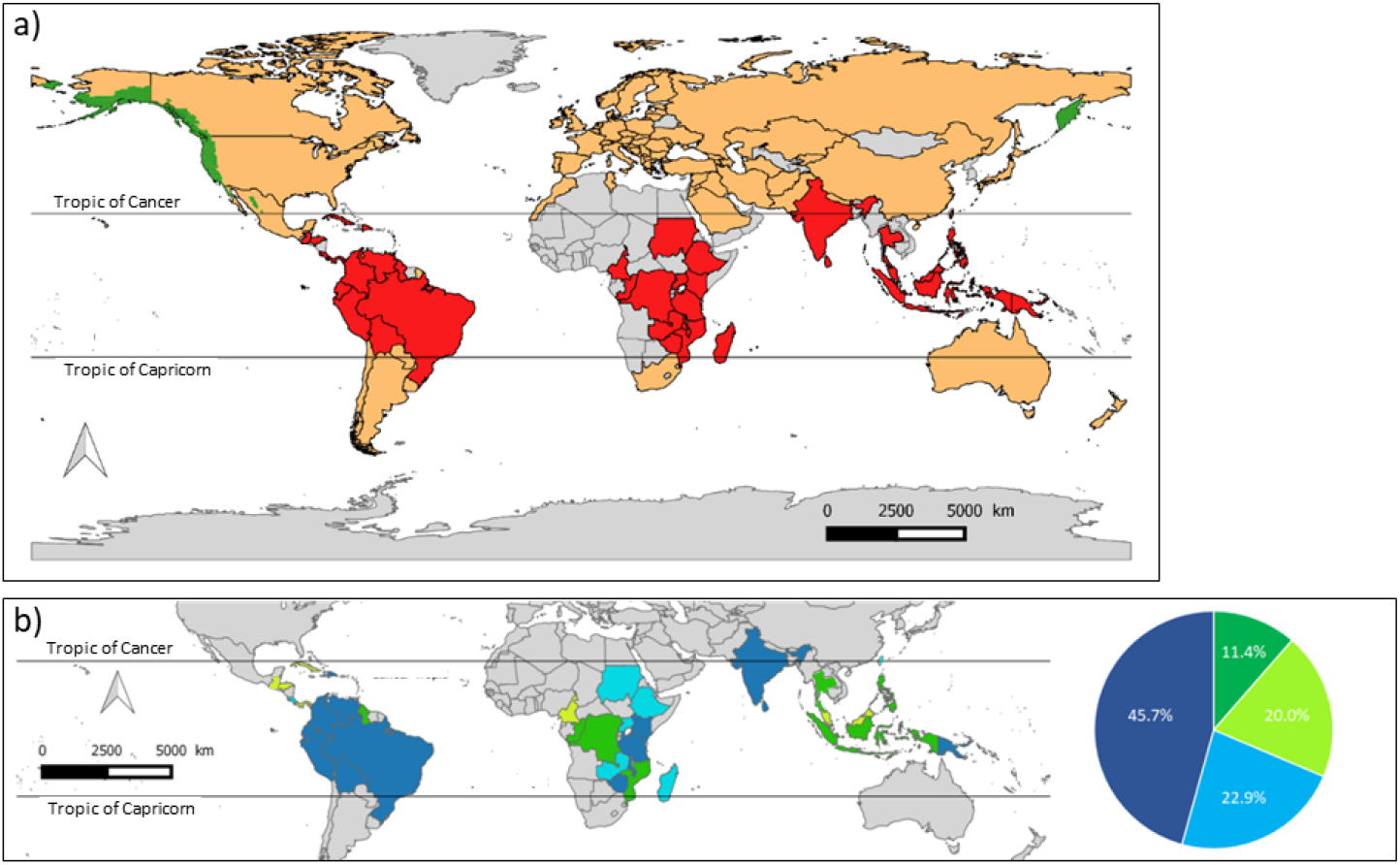
Current distribution of rainbow trout worldwide. a) Native-distribution (green), temperate (orange) and tropical (red) countries in which the species has been introduced (adapted from Crawford and Muir (2008). b) Results of introducing rainbow trout into tropical countries: evidence of the presence of self-sustaining populations in 1988 (in the review by Welcomme (1988), light blue) and after 1988 (dark blue), no information available about the presence of self-sustaining populations (light green), and no evidence of self-sustaining populations (dark green). See Table 1 for supporting information and references.

Since the middle of the 20^th^ century, rainbow trout has been massively introduced worldwide for aquaculture and recreational purposes (Welcomme 1988). According to the FAO (2003), at least 97 countries have introduced it (Fig. 1a), including 38 in the tropics (Table S1). To our knowledge, rainbow trout is the only salmonid species that lives in the tropics. Lethal temperature is the main reason that introductions have failed in Puerto Rico, the Philippines, Trinidad and Mauritius (MacCrimmon 1971). Nevertheless, the existence of self-sustaining populations (i.e. recruitment unsupported by and independent of continued human interactions, *sensu* Copp *et al*. (2005)) has been reported in nearly 70% of tropical countries where the species was introduced (Fig. 1b, Table S1). Tropical populations of rainbow trout are presumably found at high elevations, where the ambient temperature is cool (Therezien 1960; Péfaur & Sierra 1998; Ngugi & Green 2007; Vila *et al*. 2007; Kadye *et al*. 2013; Vimos *et al*. 2015), such as in the Nyanga Mountains (>1,800 m) in Zimbabwe or Lake Titicaca (3,812 m) in Peru and Bolivia. However, the dual influence of water temperature and elevation on the success of introduced rainbow trout in the tropics has not been clearly determined. In the current context of global warming, understanding how rainbow trout populations respond to high temperatures in tropical regions may help better predict the future of salmonids in their native ranges.

**Table 1.**
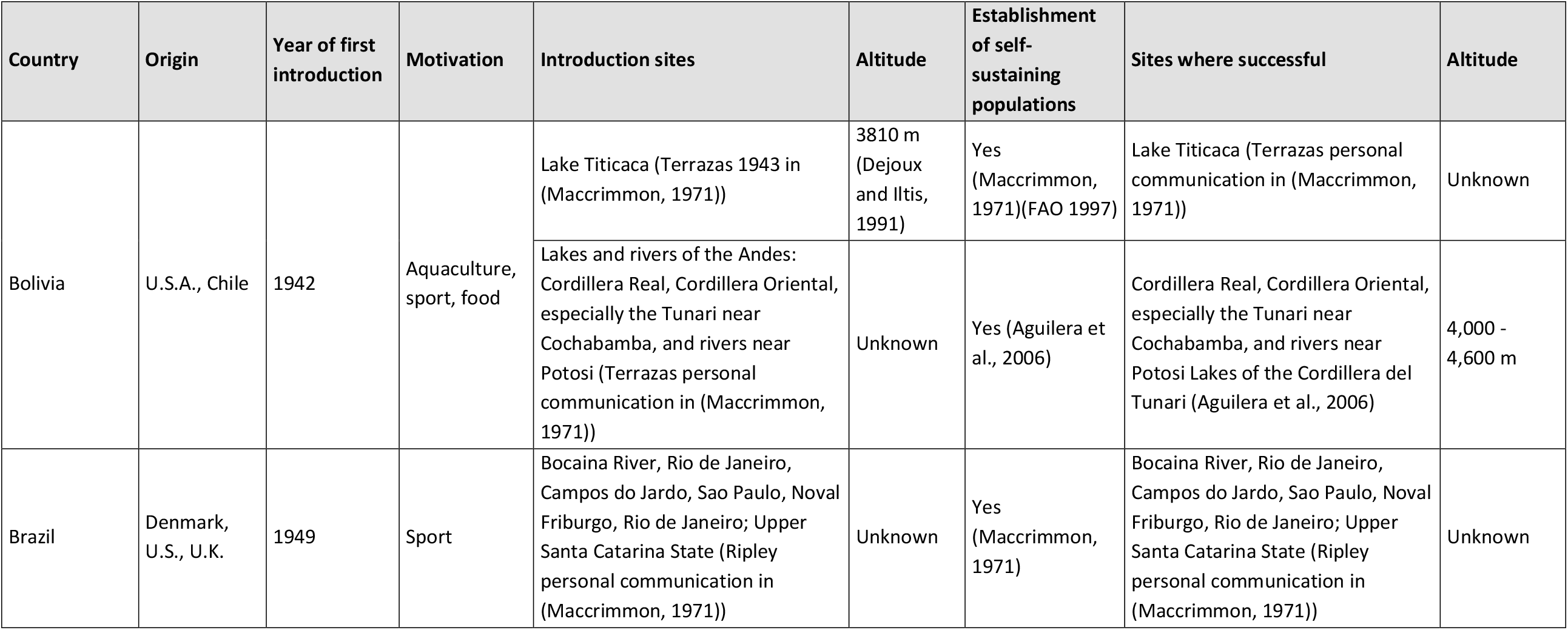
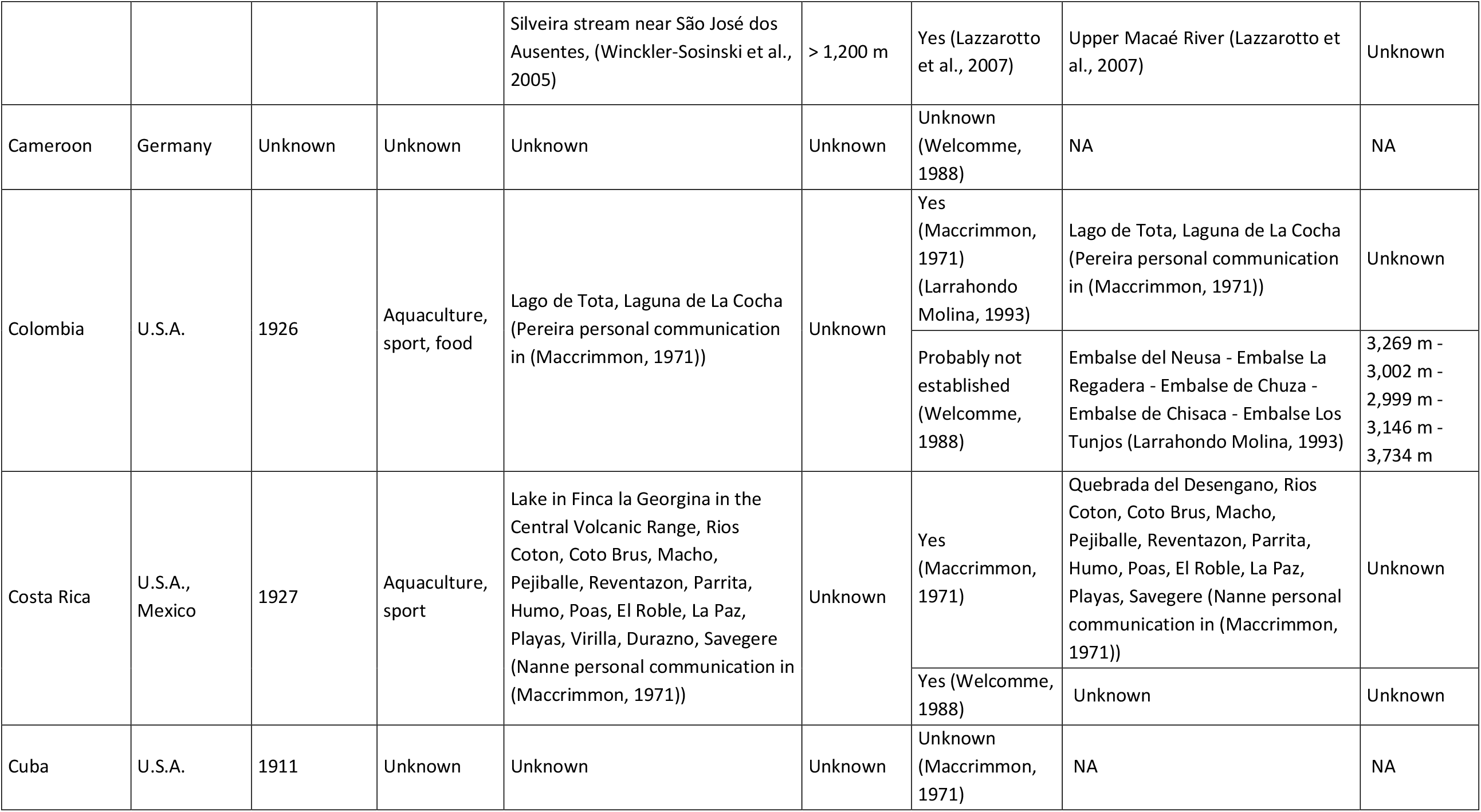
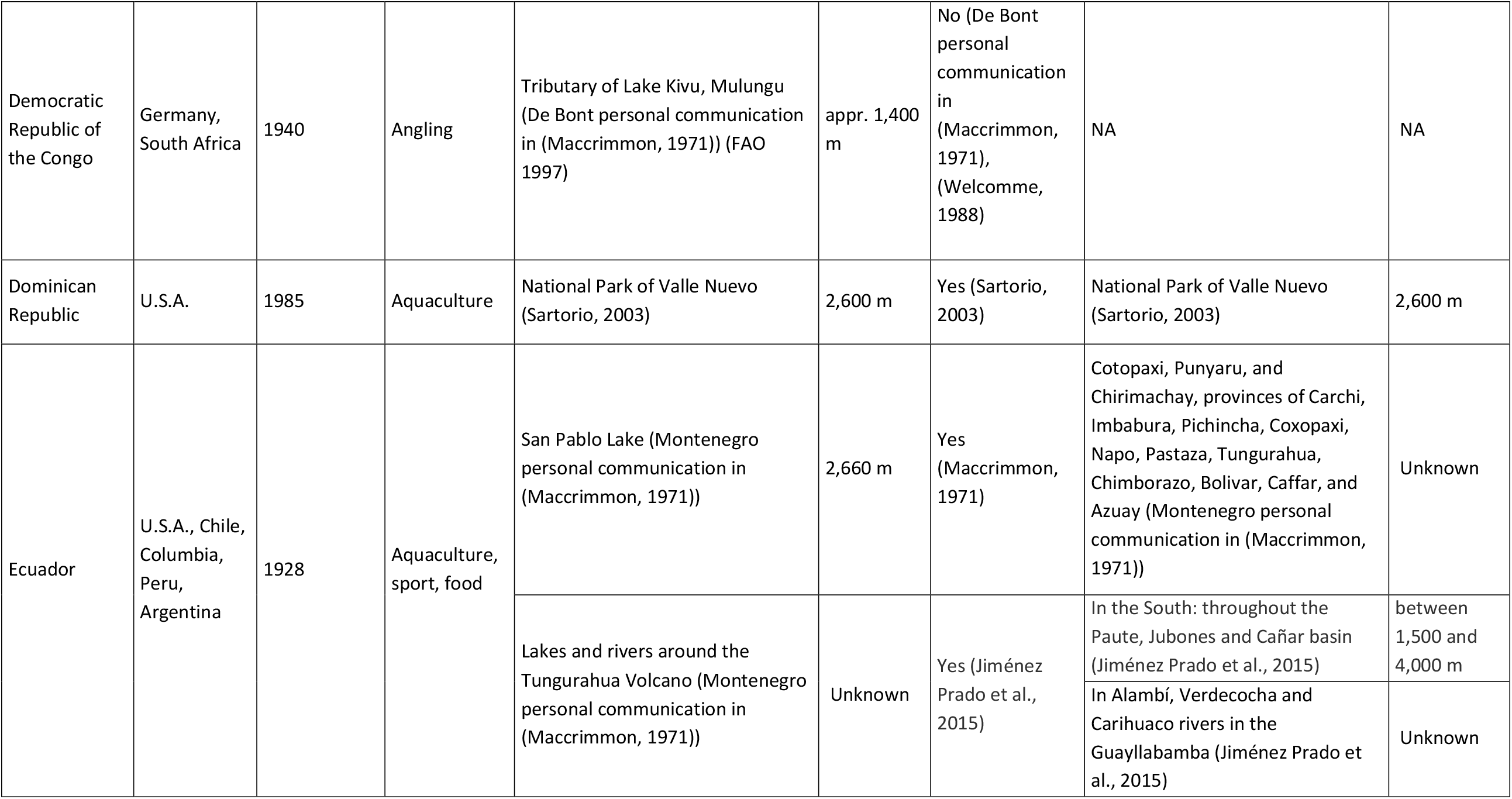
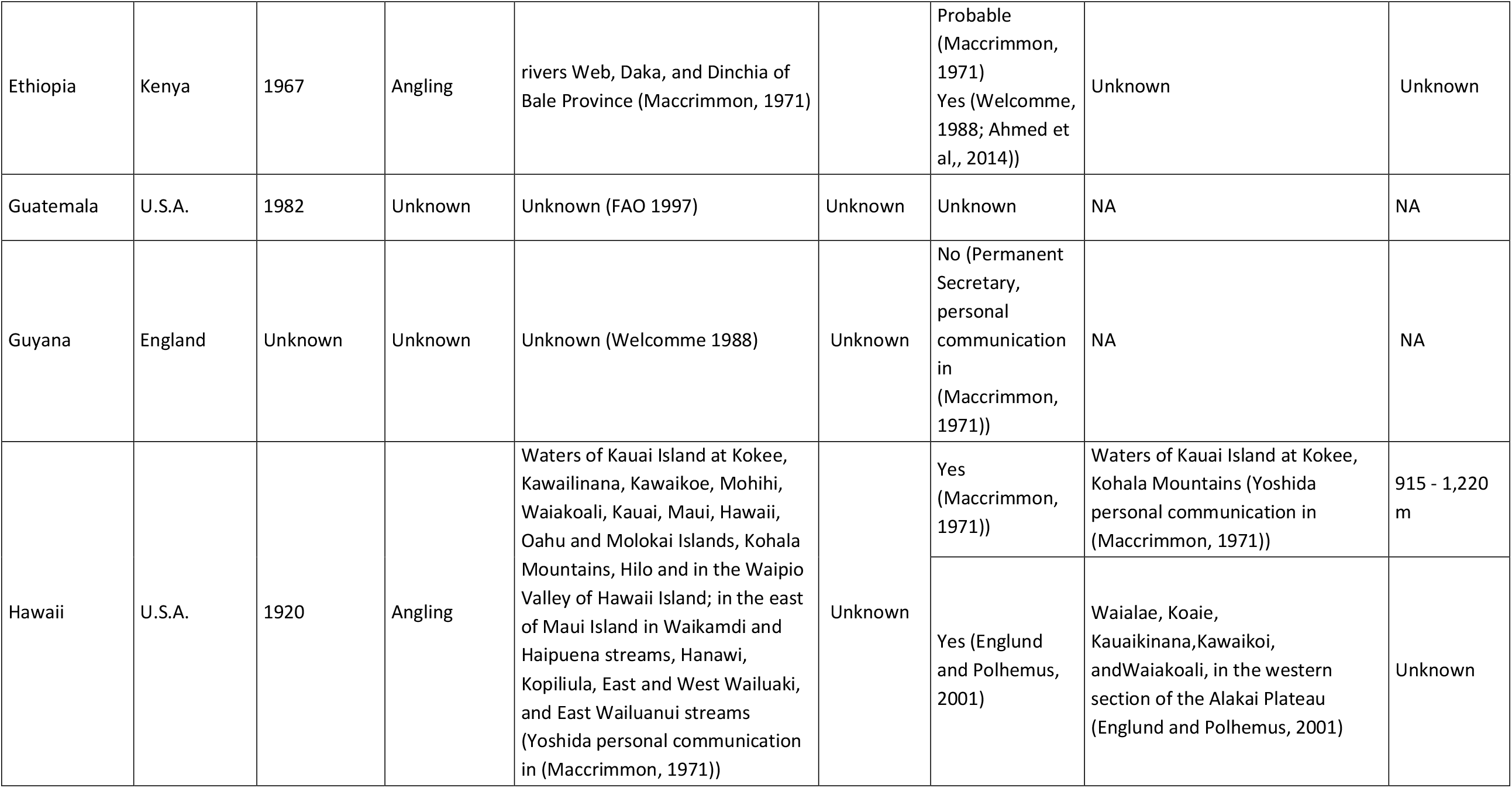
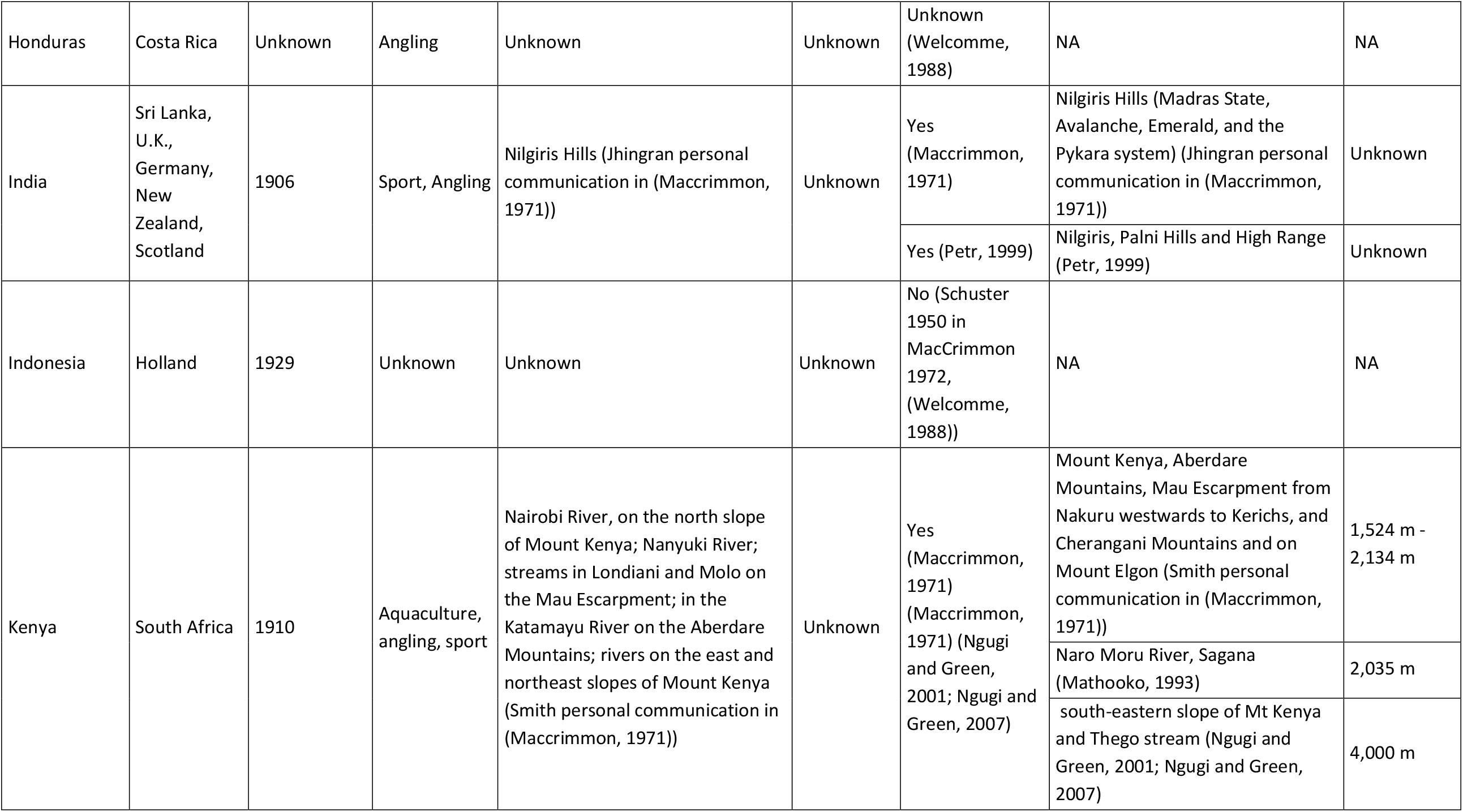
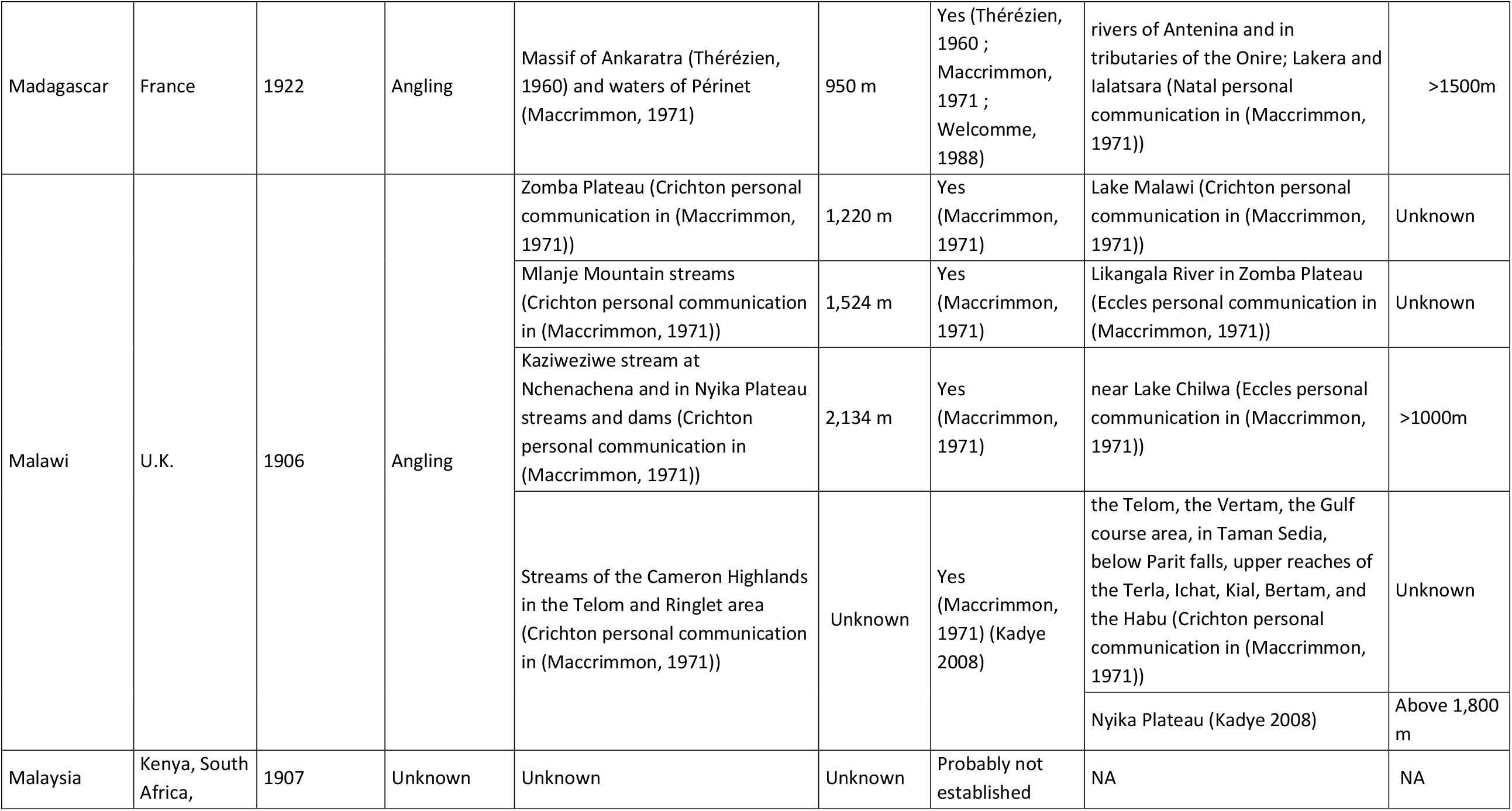
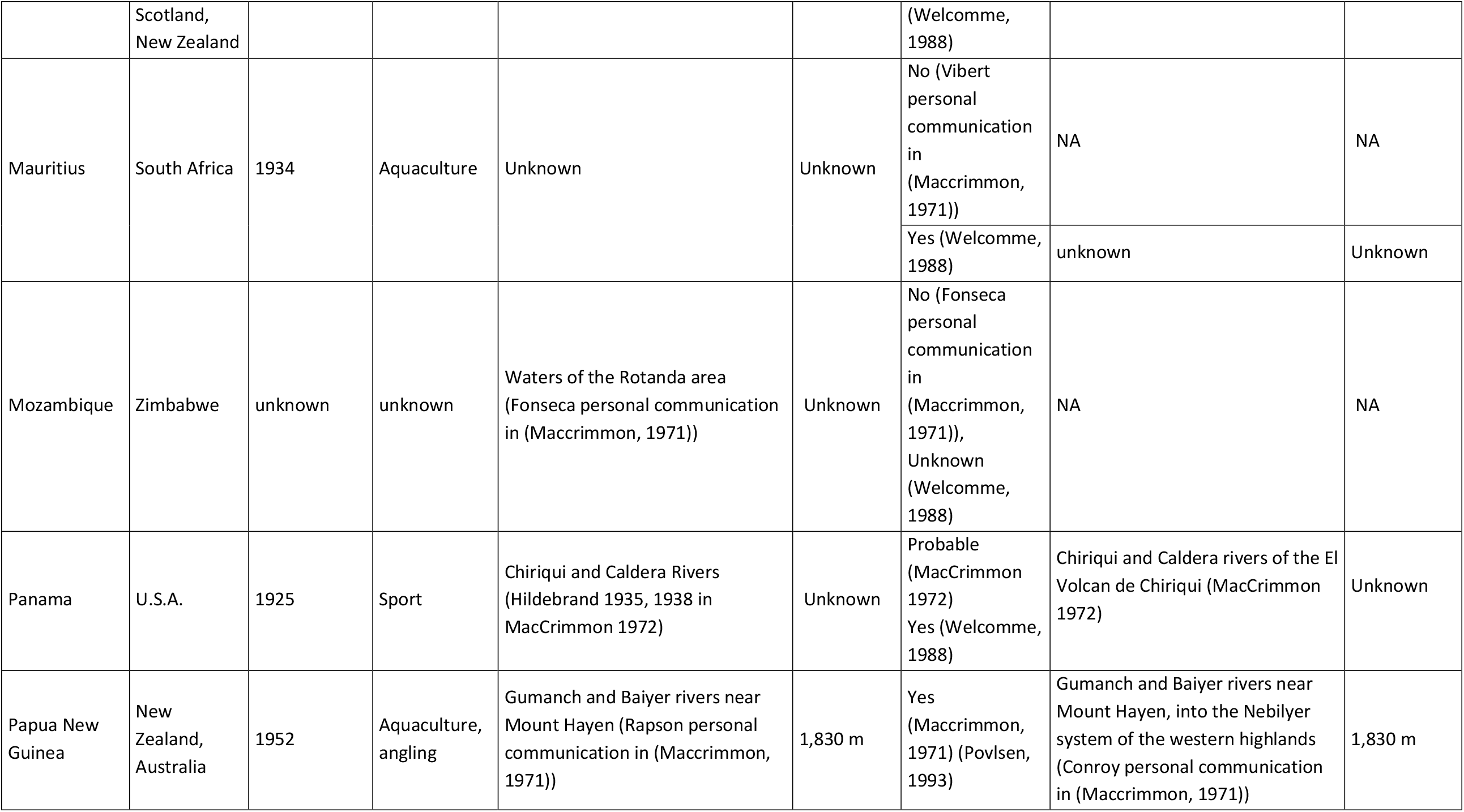
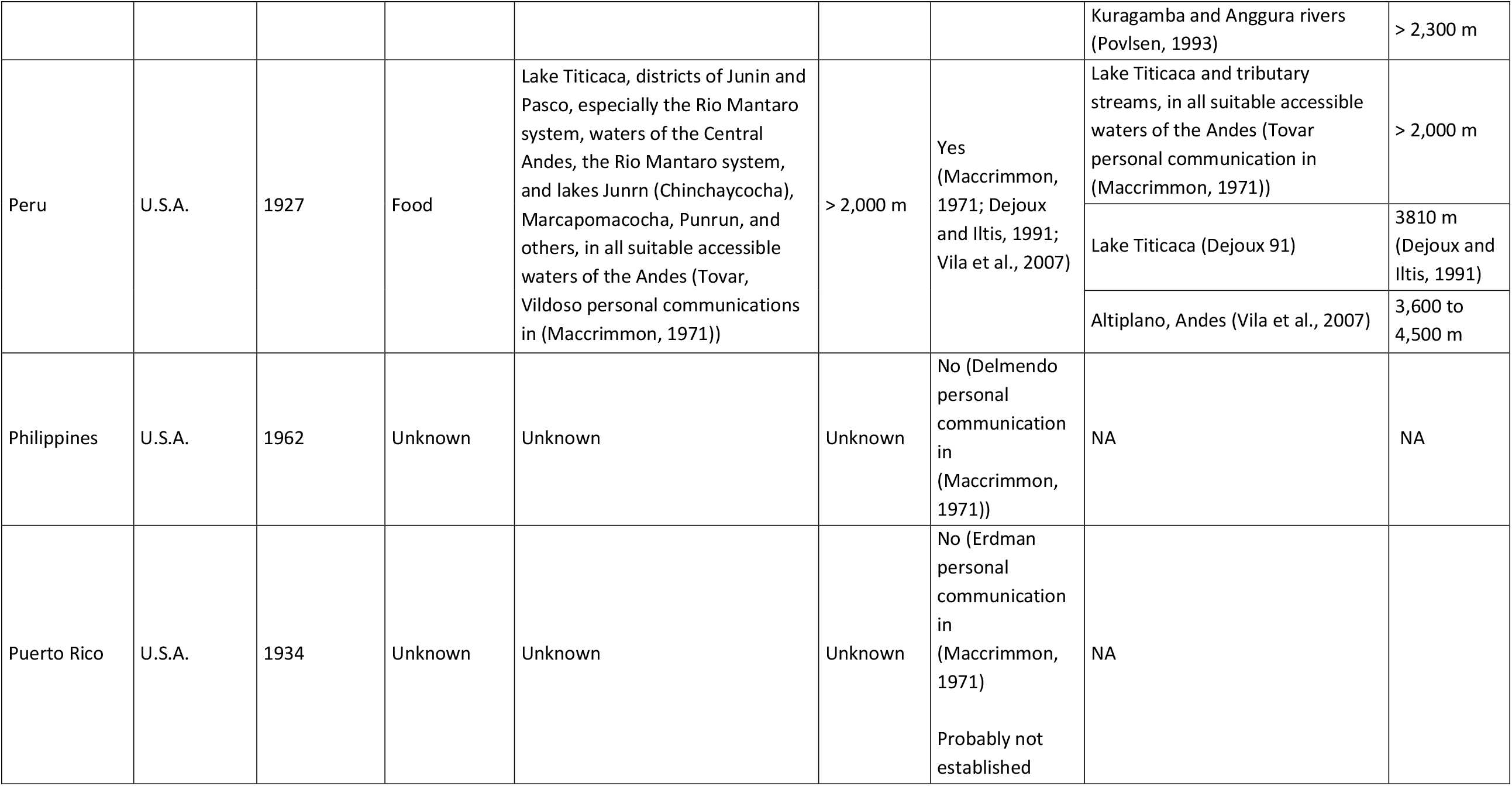
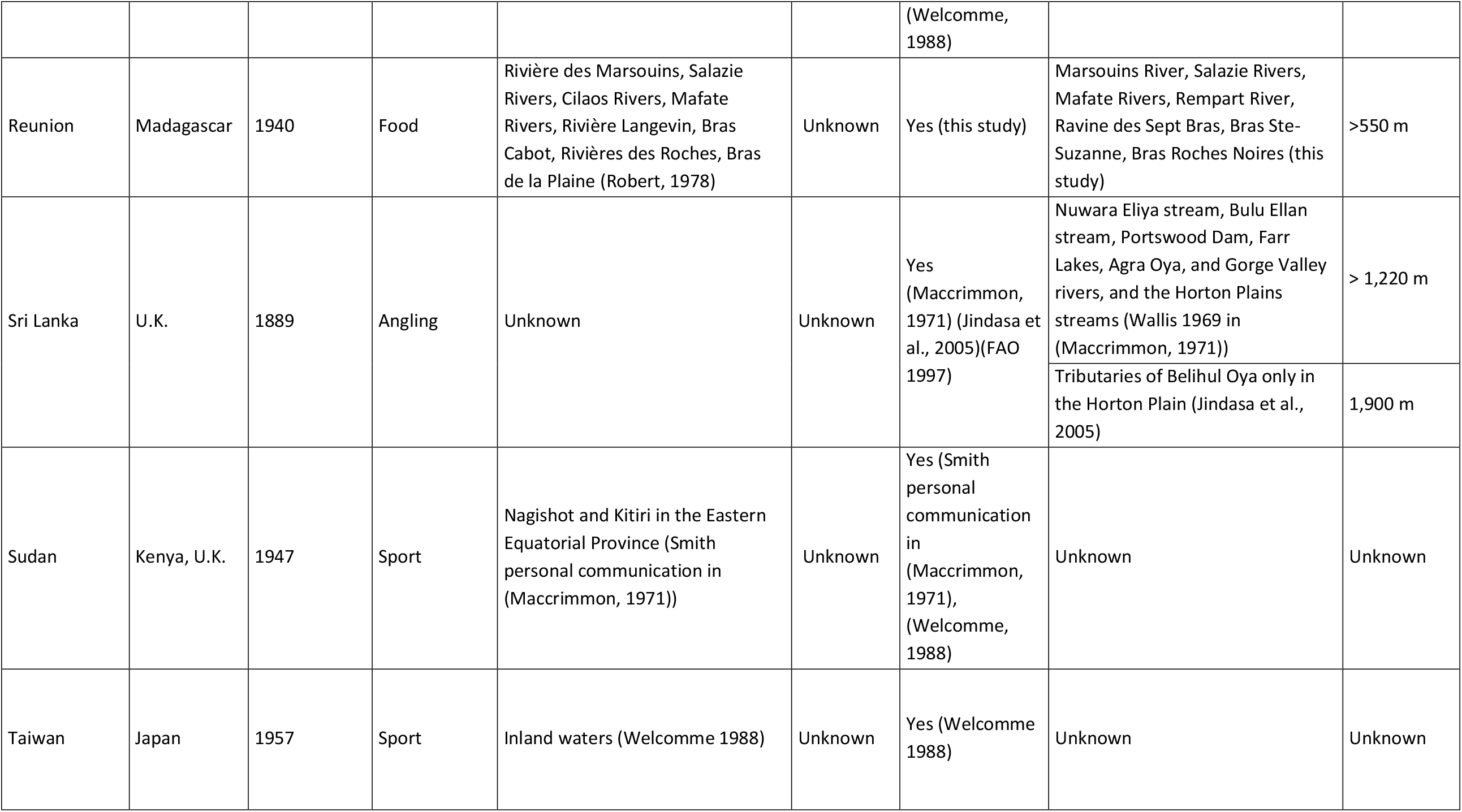
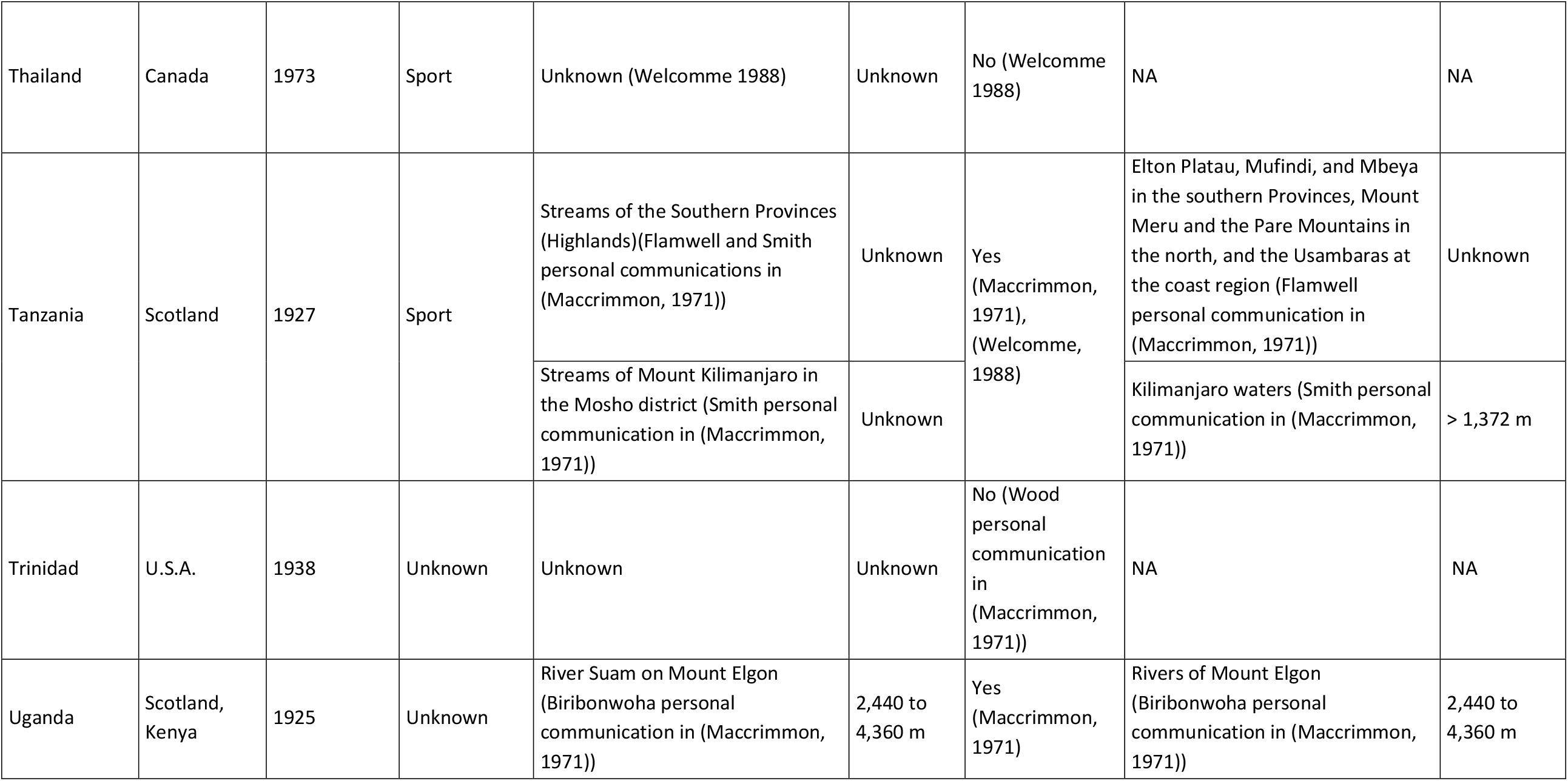
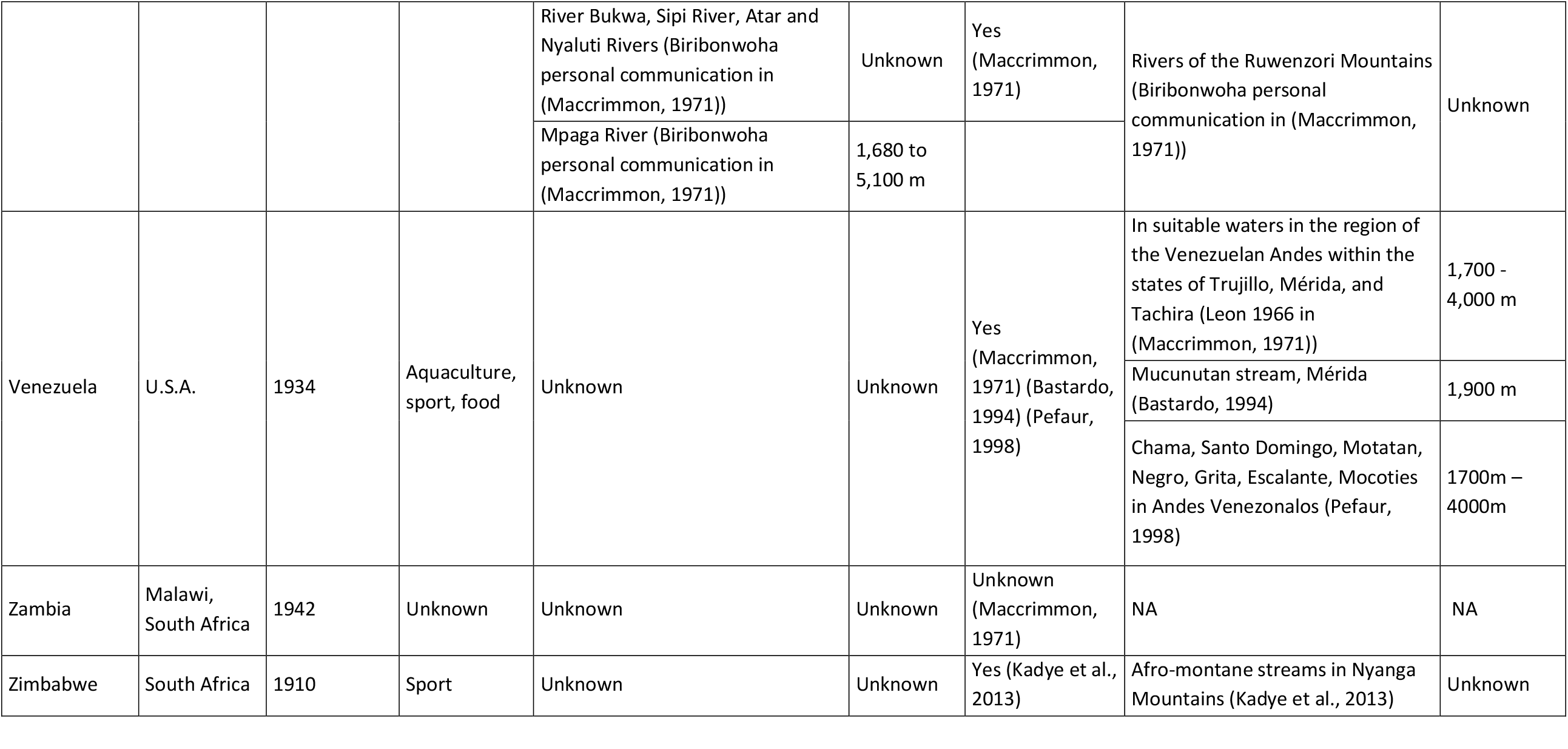
Evidence of introductions and the presence of self-sustaining populations of rainbow trout in the tropics.

| Country | Origin | Year of first introduction | Motivation | Introduction sites | Altitude | Establishment of self-sustaining populations | Sites where successful | Altitude |
| --- | --- | --- | --- | --- | --- | --- | --- | --- |
| Bolivia | U.S.A., Chile | 1942 | Aquaculture, sport, food | Lake Titicaca (Terrazas 1943 in (Maccrimmon, 1971)) | 3810 m (Dejoux and Iltis, 1991) | Yes (Maccrimmon, 1971)(FAO 1997) | Lake Titicaca (Terrazas personal communication in (Maccrimmon, 1971)) | Unknown |
|  |  |  |  | Lakes and rivers of the Andes: Cordillera Real, Cordillera Oriental, especially the Tunari near Cochabamba, and rivers near Potosi (Terrazas personal communication in (Maccrimmon, 1971)) | Unknown | Yes (Aguilera et al., 2006) | Cordillera Real, Cordillera Oriental, especially the Tunari near Cochabamba, and rivers near Potosi Lakes of the Cordillera del Tunari (Aguilera et al., 2006) | 4,000 - 4,600 m |
| Brazil | Denmark, U.S., U.K. | 1949 | Sport | Bocaina River, Rio de Janeiro, Campos do Jardo, Sao Paulo, Noval Friburgo, Rio de Janeiro; Upper Santa Catarina State (Ripley personal communication in (Maccrimmon, 1971)) | Unknown | Yes (Maccrimmon, 1971) | Bocaina River, Rio de Janeiro, Campos do Jardo, Sao Paulo, Noval Friburgo, Rio de Janeiro; Upper Santa Catarina State (Ripley personal communication in (Maccrimmon, 1971)) | Unknown |
|  |  |  |  | Silveira stream near São José dos Ausentes, (Winckler-Sosinski et al., 2005) | > 1,200 m | Yes (Lazzarotto et al., 2007) | Upper Macaé River (Lazzarotto et al., 2007) | Unknown |
| Cameroon | Germany | Unknown | Unknown | Unknown | Unknown | Unknown (Welcomme, 1988) | NA | NA |
| Colombia | U.S.A. | 1926 | Aquaculture, sport, food | Lago de Tota, Laguna de La Cocha (Pereira personal communication in (Maccrimmon, 1971)) | Unknown | Yes (Maccrimmon, 1971) (Larrahondo Molina, 1993) | Lago de Tota, Laguna de La Cocha (Pereira personal communication in (Maccrimmon, 1971)) | Unknown |
|  |  |  |  |  |  | Probably not established (Welcomme, 1988) | Embalse del Neusa - Embalse La Regadera - Embalse de Chuza - Embalse de Chisaca - Embalse Los Tunjos (Larrahondo Molina, 1993) | 3,269 m - 3,002 m - 2,999 m - 3,146 m - 3,734 m |
| Costa Rica | U.S.A., Mexico | 1927 | Aquaculture, sport | Lake in Finca la Georgina in the Central Volcanic Range, Rios Coton, Coto Brus, Macho, Pejiballe, Reventazon, Parrita, Humo, Poas, El Roble, La Paz, Playas, Virilla, Durazno, Savegere (Nanne personal communication in (Maccrimmon, 1971)) | Unknown | Yes (Maccrimmon, 1971) | Quebrada del Desengano, Rios Coton, Coto Brus, Macho, Pejiballe, Reventazon, Parrita, Humo, Poas, El Roble, La Paz, Playas, Savegere (Nanne personal communication in (Maccrimmon, 1971)) | Unknown |
|  |  |  |  |  |  | Yes (Welcomme, 1988) | Unknown | Unknown |
| Cuba | U.S.A. | 1911 | Unknown | Unknown | Unknown | Unknown (Maccrimmon, 1971) | NA | NA |
| Democratic Republic of the Congo | Germany, South Africa | 1940 | Angling | Tributary of Lake Kivu, Mulungu (De Bont personal communication in (Maccrimmon, 1971)) (FAO 1997) | appr. 1,400 m | No (De Bont personal communication in (Maccrimmon, 1971), (Welcomme, 1988) | NA | NA |
| Dominican Republic | U.S.A. | 1985 | Aquaculture | National Park of Valle Nuevo (Sartorio, 2003) | 2,600 m | Yes (Sartorio, 2003) | National Park of Valle Nuevo (Sartorio, 2003) | 2,600 m |
| Ecuador | U.S.A., Chile, Columbia, Peru, Argentina | 1928 | Aquaculture, sport, food | San Pablo Lake (Montenegro personal communication in (Maccrimmon, 1971)) | 2,660 m | Yes (Maccrimmon, 1971) | Cotopaxi, Punyaru, and Chirimachay, provinces of Carchi, Imbabura, Pichincha, Coxopaxi, Napo, Pastaza, Tungurahua, Chimborazo, Bolivar, Caffar, and Azuay (Montenegro personal communication in (Maccrimmon, 1971)) | Unknown |
|  |  |  |  | Lakes and rivers around the Tungurahua Volcano (Montenegro personal communication in (Maccrimmon, 1971)) | Unknown | Yes (Jiménez Prado et al., 2015) | In the South: throughout the Paute, Jubones and Cañar basin (Jiménez Prado et al., 2015) | between 1,500 and 4,000 m |
|  |  |  |  |  |  |  | In Alambí, Verdecocha and Carihuaco rivers in the Guayllabamba (Jiménez Prado et al., 2015) | Unknown |
| Ethiopia | Kenya | 1967 | Angling | rivers Web, Daka, and Dinchia of Bale Province (Maccrimmon, 1971) |  | Probable (Maccrimmon, 1971)<br>Yes (Welcomme, 1988; Ahmed et al., 2014)) | Unknown | Unknown |
| Guatemala | U.S.A. | 1982 | Unknown | Unknown (FAO 1997) | Unknown | Unknown | NA | NA |
| Guyana | England | Unknown | Unknown | Unknown (Welcomme 1988) | Unknown | No (Permanent Secretary, personal communication in (Maccrimmon, 1971)) | NA | NA |
| Hawaii | U.S.A. | 1920 | Angling | Waters of Kauai Island at Kokee, Kawailinana, Kawaikoe, Mohihi, Waiakoali, Kauai, Maui, Hawaii, Oahu and Molokai Islands, Kohala Mountains, Hilo and in the Waipio Valley of Hawaii Island; in the east of Maui Island in Waikamdi and Haipuena streams, Hanawi, Kopiliula, East and West Wailuaki, and East Wailuanui streams (Yoshida personal communication in (Maccrimmon, 1971)) | Unknown | Yes (Maccrimmon, 1971)) | Waters of Kauai Island at Kokee, Kohala Mountains (Yoshida personal communication in (Maccrimmon, 1971)) | 915 - 1,220 m |
|  |  |  |  |  |  | Yes (Englund and Polhemus, 2001) | Waialae, Koaie, Kauaikinana, Kawaikoi, and Waiakoali, in the western section of the Alakai Plateau (Englund and Polhemus, 2001) | Unknown |
| Honduras | Costa Rica | Unknown | Angling | Unknown | Unknown | Unknown (Welcomme, 1988) | NA | NA |
| India | Sri Lanka, U.K., Germany, New Zealand, Scotland | 1906 | Sport, Angling | Nilgiris Hills (Jhingran personal communication in (Maccrimmon, 1971)) | Unknown | Yes (Maccrimmon, 1971) | Nilgiris Hills (Madras State, Avalanche, Emerald, and the Pykara system) (Jhingran personal communication in (Maccrimmon, 1971)) | Unknown |
|  |  |  |  |  |  | Yes (Petr, 1999) | Nilgiris, Palni Hills and High Range (Petr, 1999) | Unknown |
| Indonesia | Holland | 1929 | Unknown | Unknown | Unknown | No (Schuster 1950 in MacCrimmon 1972, (Welcomme, 1988)) | NA | NA |
| Kenya | South Africa | 1910 | Aquaculture, angling, sport | Nairobi River, on the north slope of Mount Kenya; Nanyuki River; streams in Londiani and Molo on the Mau Escarpment; in the Katamayu River on the Aberdare Mountains; rivers on the east and northeast slopes of Mount Kenya (Smith personal communication in (Maccrimmon, 1971)) | Unknown | Yes (Maccrimmon, 1971) (Maccrimmon, 1971) (Ngugi and Green, 2001; Ngugi and Green, 2007) | Mount Kenya, Aberdare Mountains, Mau Escarpment from Nakuru westwards to Kerichs, and Cherangani Mountains and on Mount Elgon (Smith personal communication in (Maccrimmon, 1971)) | 1,524 m - 2,134 m |
|  |  |  |  |  |  |  | Naro Moru River, Sagana (Mathooko, 1993) | 2,035 m |
|  |  |  |  |  |  |  | south-eastern slope of Mt Kenya and Thego stream (Ngugi and Green, 2001; Ngugi and Green, 2007) | 4,000 m |
| Madagascar | France | 1922 | Angling | Massif of Ankaratra (Thérézien, 1960) and waters of Périnet (Maccrimmon, 1971) | 950 m | Yes (Thérézien, 1960 ; Maccrimmon, 1971 ; Welcomme, 1988) | rivers of Antenina and in tributaries of the Onire; Lakera and lalatsara (Natal personal communication in (Maccrimmon, 1971)) | >1500m |
| Malawi | U.K. | 1906 | Angling | Zomba Plateau (Crichton personal communication in (Maccrimmon, 1971)) | 1,220 m | Yes (Maccrimmon, 1971) | Lake Malawi (Crichton personal communication in (Maccrimmon, 1971)) | Unknown |
|  |  |  |  | Mlanje Mountain streams (Crichton personal communication in (Maccrimmon, 1971)) | 1,524 m | Yes (Maccrimmon, 1971) | Likangala River in Zomba Plateau (Eccles personal communication in (Maccrimmon, 1971)) | Unknown |
|  |  |  |  | Kaziweziwe stream at Nchenachena and in Nyika Plateau streams and dams (Crichton personal communication in (Maccrimmon, 1971)) | 2,134 m | Yes (Maccrimmon, 1971) | near Lake Chilwa (Eccles personal communication in (Maccrimmon, 1971)) | >1000m |
|  |  |  |  | Streams of the Cameron Highlands in the Telom and Ringlet area (Crichton personal communication in (Maccrimmon, 1971)) | Unknown | Yes (Maccrimmon, 1971) (Kadye 2008) | the Telom, the Vertam, the Gulf course area, in Taman Sedia, below Parit falls, upper reaches of the Terla, Ichat, Kial, Bertam, and the Habu (Crichton personal communication in (Maccrimmon, 1971)) | Unknown |
|  |  |  |  |  |  |  | Nyika Plateau (Kadye 2008) | Above 1,800 m |
| Malaysia | Kenya, South Africa, | 1907 | Unknown | Unknown | Unknown | Probably not established | NA | NA |
|  | Scotland,<br>New Zealand |  |  |  |  | (Welcomme,<br>1988) |  |  |
| Mauritius | South Africa | 1934 | Aquaculture | Unknown | Unknown | No (Vibert<br>personal<br>communication<br>in<br>(Maccrimmon,<br>1971)) | NA | NA |
|  |  |  |  |  |  | Yes (Welcomme,<br>1988) | unknown | Unknown |
| Mozambique | Zimbabwe | unknown | unknown | Waters of the Rotanda area<br>(Fonseca personal communication<br>in (Maccrimmon, 1971)) | Unknown | No (Fonseca<br>personal<br>communication<br>in<br>(Maccrimmon,<br>1971)),<br>Unknown<br>(Welcomme,<br>1988) | NA | NA |
| Panama | U.S.A. | 1925 | Sport | Chiriqui and Caldera Rivers<br>(Hildebrand 1935, 1938 in<br>MacCrimmon 1972) | Unknown | Probable<br>(MacCrimmon<br>1972)<br>Yes (Welcomme,<br>1988) | Chiriqui and Caldera rivers of the El<br>Volcan de Chiriqui (MacCrimmon<br>1972) | Unknown |
| Papua New<br>Guinea | New<br>Zealand,<br>Australia | 1952 | Aquaculture,<br>angling | Gumanch and Baiyer rivers near<br>Mount Hayen (Rapson personal<br>communication in (Maccrimmon,<br>1971)) | 1,830 m | Yes<br>(Maccrimmon,<br>1971) (Povlsen,<br>1993) | Gumanch and Baiyer rivers near<br>Mount Hayen, into the Nebilyer<br>system of the western highlands<br>(Conroy personal communication<br>in (Maccrimmon, 1971)) | 1,830 m |
|  |  |  |  |  |  |  | Kuragamba and Anggura rivers (Povlsen, 1993) | > 2,300 m |
| Peru | U.S.A. | 1927 | Food | Lake Titicaca, districts of Junin and Pasco, especially the Rio Mantaro system, waters of the Central Andes, the Rio Mantaro system, and lakes Junrn (Chinchaycocha), Marcapomacocha, Punrun, and others, in all suitable accessible waters of the Andes (Tovar, Vildoso personal communications in (Maccrimmon, 1971)) | > 2,000 m | Yes (Maccrimmon, 1971; Dejoux and Iltis, 1991; Vila et al., 2007) | Lake Titicaca and tributary streams, in all suitable accessible waters of the Andes (Tovar personal communication in (Maccrimmon, 1971)) | > 2,000 m |
|  |  |  |  |  |  |  | Lake Titicaca (Dejoux 91) | 3810 m (Dejoux and Iltis, 1991) |
|  |  |  |  |  |  |  | Altiplano, Andes (Vila et al., 2007) | 3,600 to 4,500 m |
| Philippines | U.S.A. | 1962 | Unknown | Unknown | Unknown | No (Delmendo personal communication in (Maccrimmon, 1971)) | NA | NA |
| Puerto Rico | U.S.A. | 1934 | Unknown | Unknown | Unknown | No (Erdman personal communication in (Maccrimmon, 1971))<br>Probably not established | NA |  |
|  |  |  |  |  |  | (Welcomme, 1988) |  |  |
| Reunion | Madagascar | 1940 | Food | Rivière des Marsouins, Salazie Rivers, Cilaos Rivers, Mafate Rivers, Rivière Langevin, Bras Cabot, Rivières des Roches, Bras de la Plaine (Robert, 1978) | Unknown | Yes (this study) | Marsouins River, Salazie Rivers, Mafate Rivers, Rempart River, Ravine des Sept Bras, Bras Ste-Suzanne, Bras Roches Noires (this study) | >550 m |
| Sri Lanka | U.K. | 1889 | Angling | Unknown | Unknown | Yes (Maccrimmon, 1971) (Jindasa et al., 2005)(FAO 1997) | Nuwara Eliya stream, Bulu Ellan stream, Portswood Dam, Farr Lakes, Agra Oya, and Gorge Valley rivers, and the Horton Plains streams (Wallis 1969 in (Maccrimmon, 1971)) | > 1,220 m |
|  |  |  |  |  |  |  | Tributaries of Belihul Oya only in the Horton Plain (Jindasa et al., 2005) | 1,900 m |
| Sudan | Kenya, U.K. | 1947 | Sport | Nagishot and Kitiri in the Eastern Equatorial Province (Smith personal communication in (Maccrimmon, 1971)) | Unknown | Yes (Smith personal communication in (Maccrimmon, 1971), (Welcomme, 1988) | Unknown | Unknown |
| Taiwan | Japan | 1957 | Sport | Inland waters (Welcomme 1988) | Unknown | Yes (Welcomme 1988) | Unknown | Unknown |
| Thailand | Canada | 1973 | Sport | Unknown (Welcomme 1988) | Unknown | No (Welcomme 1988) | NA | NA |
| Tanzania | Scotland | 1927 | Sport | Streams of the Southern Provinces (Highlands)(Flamwell and Smith personal communications in (Maccrimmon, 1971)) | Unknown | Yes (Maccrimmon, 1971), (Welcomme, 1988) | Elton Platau, Mufindi, and Mbeya in the southern Provinces, Mount Meru and the Pare Mountains in the north, and the Usambaras at the coast region (Flamwell personal communication in (Maccrimmon, 1971)) | Unknown |
|  |  |  |  | Streams of Mount Kilimanjaro in the Mosho district (Smith personal communication in (Maccrimmon, 1971)) | Unknown |  | Kilimanjaro waters (Smith personal communication in (Maccrimmon, 1971)) | > 1,372 m |
| Trinidad | U.S.A. | 1938 | Unknown | Unknown | Unknown | No (Wood personal communication in (Maccrimmon, 1971)) | NA | NA |
| Uganda | Scotland, Kenya | 1925 | Unknown | River Suam on Mount Elgon (Biribonwoha personal communication in (Maccrimmon, 1971)) | 2,440 to 4,360 m | Yes (Maccrimmon, 1971) | Rivers of Mount Elgon (Biribonwoha personal communication in (Maccrimmon, 1971)) | 2,440 to 4,360 m |
|  |  |  |  | River Bukwa, Sipi River, Atar and Nyaluti Rivers (Biribonwoha personal communication in (Maccrimmon, 1971)) | Unknown | Yes (Maccrimmon, 1971) | Rivers of the Ruwenzori Mountains (Biribonwoha personal communication in (Maccrimmon, 1971)) | Unknown |
|  |  |  |  | Mpaga River (Biribonwoha personal communication in (Maccrimmon, 1971)) | 1,680 to 5,100 m |  |  |  |
| Venezuela | U.S.A. | 1934 | Aquaculture, sport, food | Unknown | Unknown | Yes (Maccrimmon, 1971) (Bastardo, 1994) (Pefaur, 1998) | In suitable waters in the region of the Venezuelan Andes within the states of Trujillo, Mérida, and Tachira (Leon 1966 in (Maccrimmon, 1971)) | 1,700 - 4,000 m |
|  |  |  |  |  |  |  | Mucunutan stream, Mérida (Bastardo, 1994) | 1,900 m |
|  |  |  |  |  |  |  | Chama, Santo Domingo, Motatan, Negro, Grita, Escalante, Mocoties in Andes Venezonalos (Pefaur, 1998) | 1700m – 4000m |
| Zambia | Malawi, South Africa | 1942 | Unknown | Unknown | Unknown | Unknown (Maccrimmon, 1971) | NA | NA |
| Zimbabwe | South Africa | 1910 | Sport | Unknown | Unknown | Yes (Kadye et al., 2013) | Afro-montane streams in Nyanga Mountains (Kadye et al., 2013) | Unknown |

The objective of this study was to investigate how introduction effort and temperature influence the presence of self-sustaining populations of rainbow trout in tropical regions. After reviewing the literature on tropical populations of the species, we focused on the unique case study of Reunion Island (West Indian Ocean), where rainbow trout has been repeatedly introduced since 1940, with varying success. A dense network of rivers that flow from high-elevation volcanic mountains provides a large gradient of water temperatures. Since early life stages of salmonid species are particularly sensitive to high temperatures (Sullivan *et al*. 2000; Benjamin *et al*. 2013), we explored the influence of altitude, as a proxy of temperature as the main driver of successful population establishment after introduction. Warming summer temperatures are identified as a main threat for the viability of rainbow trout populations in its native range in the northwestern United States (Sloat & Osterback 2013; Brewitt & Danner 2014). In the tropics, where summer temperatures are already high, warming winters could reach a critical level that is incompatible with trout reproduction and thus become relatively more limiting than in a temperate climate. We reviewed and collected the stocking history of rainbow trout on Reunion Island to determine whether the introduction effort influenced the presence/absence of trout.

## MATERIALS & METHODS

### Study site

Reunion Island is a 2,512 km² French overseas department located in the Indian Ocean (21.2°S, 55.6°E). The highest peak of this volcanic island reaches 3,069 m above sea level. Its hydrology is characterized by heavy rain during the cyclone season (December to April). The topography and aquifers generate high spatial and temporal heterogeneity in the quantity of water in rivers (Robert 1985). The island is drained by 13 perennial rivers and 43 transient watersheds characterized by their high degree of natural habitat fragmentation and their torrentiality, which can reach up to 30,000 l/s/km² on a few km² of watershed (Robert 1985). Anthropogenic barriers, which are less common than natural fragmentation, are restricted mainly to lowland areas. The rivers of Reunion Island host 26 fish and 11 decapod crustacean species, of which 20 and 10, respectively, are indigenous catadromous species (Keith 2002). Of the seven species that were introduced to the island, only four established self-sustaining populations: tilapia (*Oreochromis niloticus*), guppy (*Poecilia reticulata*), swordtail (*Xiphophorus hellerii*) and the non-anadromous form of rainbow trout (Keith 2002).

### Introduction history of rainbow trout

The rainbow trout introduced to Reunion Island have come from Madagascar (Robert, 1975), with the initial objective to provide a new source of food for the French colony (Robert, 1978). We reviewed the available reports to update the history of trout stocking on Reunion Island. The main sources of information were publications by R. Robert (Robert 1975, 1976, 1977, 1978) and reports by the National Forest Office and the fishing association.

Despite our efforts to identify all introduction events from 1940-2019, some information from 1940-1950 and 1978-1990 may have been missing due to the loss of archival records. The presence of juvenile rainbow trout in the absence of introduction, which indicates wild reproduction, is the criterion used to define a self-sustaining population. The presence of self-sustaining rainbow trout populations is assessed by visual observation (e.g. presence of juveniles, acts of spawning, redds) and electrofishing surveys conducted by the local association for fishing and the protection of aquatic environments (hereafter, fishing association). The presence or absence of a self-sustaining population from 2016-2020 was assessed at each site where trout had been introduced, all sites where its presence was suspected. A site, the smallest spatial unit considered, was defined as a stretch of river a few hundred meters long with homogeneous habitat that is usually delineated by natural discontinuities (waterfalls). We assumed that each site’s characteristics and dynamics did not change over the five years of the study. Each site was described by a unique identification number, the river and local name, elevation (Table 2).

**Table 2.**
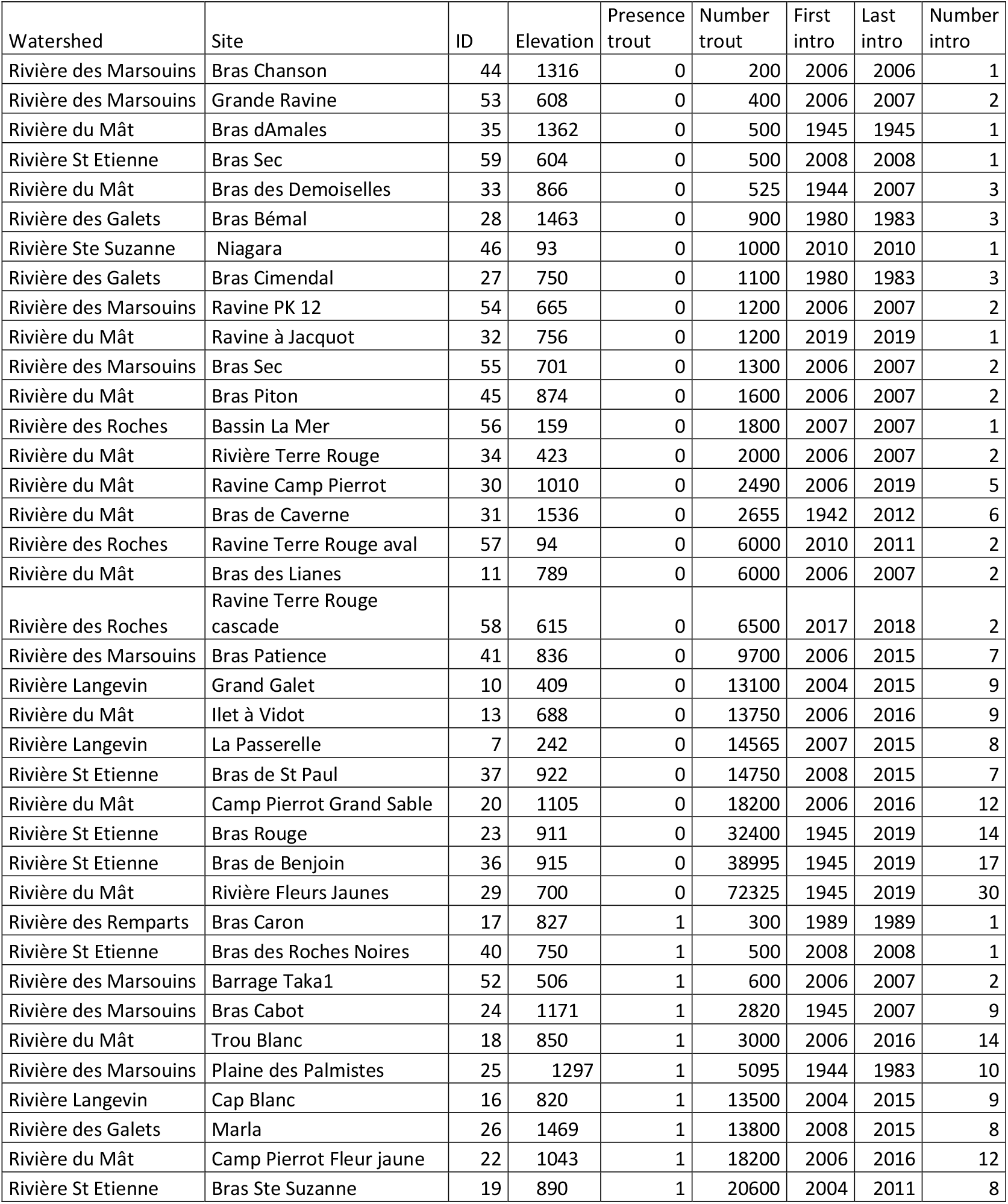
Presentation of the study sites in La Reunion Island: elevation, presence of self-sustaining population of trout, trout introduction history (number of introduction, number of trout introduced, period of introduction).

### Relating temperature to elevation

Autonomous temperature loggers provided detailed measurements of water temperature at the study sites. Water temperature was recorded daily from 2010-2021 by autonomous data loggers or fixed meteorological stations (Table 2). The local water agency (http://www.naiades.eaufrance.fr) and the fishing association provided the water temperature data. Mean monthly temperature at 12 p.m. was used as an explanatory variable in the analysis.

However, temporal and spatial coverage was often limited by the loss of loggers due to cyclones or vandalism, by the cost of the devices or by the logistical difficulties involved in deploying scientific equipment in a remote area. Since temperature depends on elevation, we explored the linear relationship between mean monthly water temperature and season/elevation using a GLM with a normal distribution of errors. The model was based on temperature and elevation data from all 33 sites (data available at https://data.inrae.fr/dataset.xhtml?persistentId=doi:10.57745/9QCPCV).

## RESULTS

### Introduction history of rainbow trout

We found evidence that rainbow trout were introduced into 8 rivers (i.e. Mât, Roches, Marsouins, Galets, Langevin, Remparts, St Etienne, Ste Suzanne) out of the 13 perennial rivers on Reunion Island (Table 2, Fig. 2). Historical records documented that 340,070 juvenile rainbow trout were released at 38 sites in these rivers or their tributaries from 1940-2019 (Table 2). The introduction effort varied greatly among sites (Fig. 2), with a single introduction of 200 trout in 2006 at site 44, but 72,325 trout introduced over more than 30 events at site 29. The presence of a self-sustaining population of trout was recorded at 10 sites of the 38 introduction sites (Table 2).

**Fig. 2.**
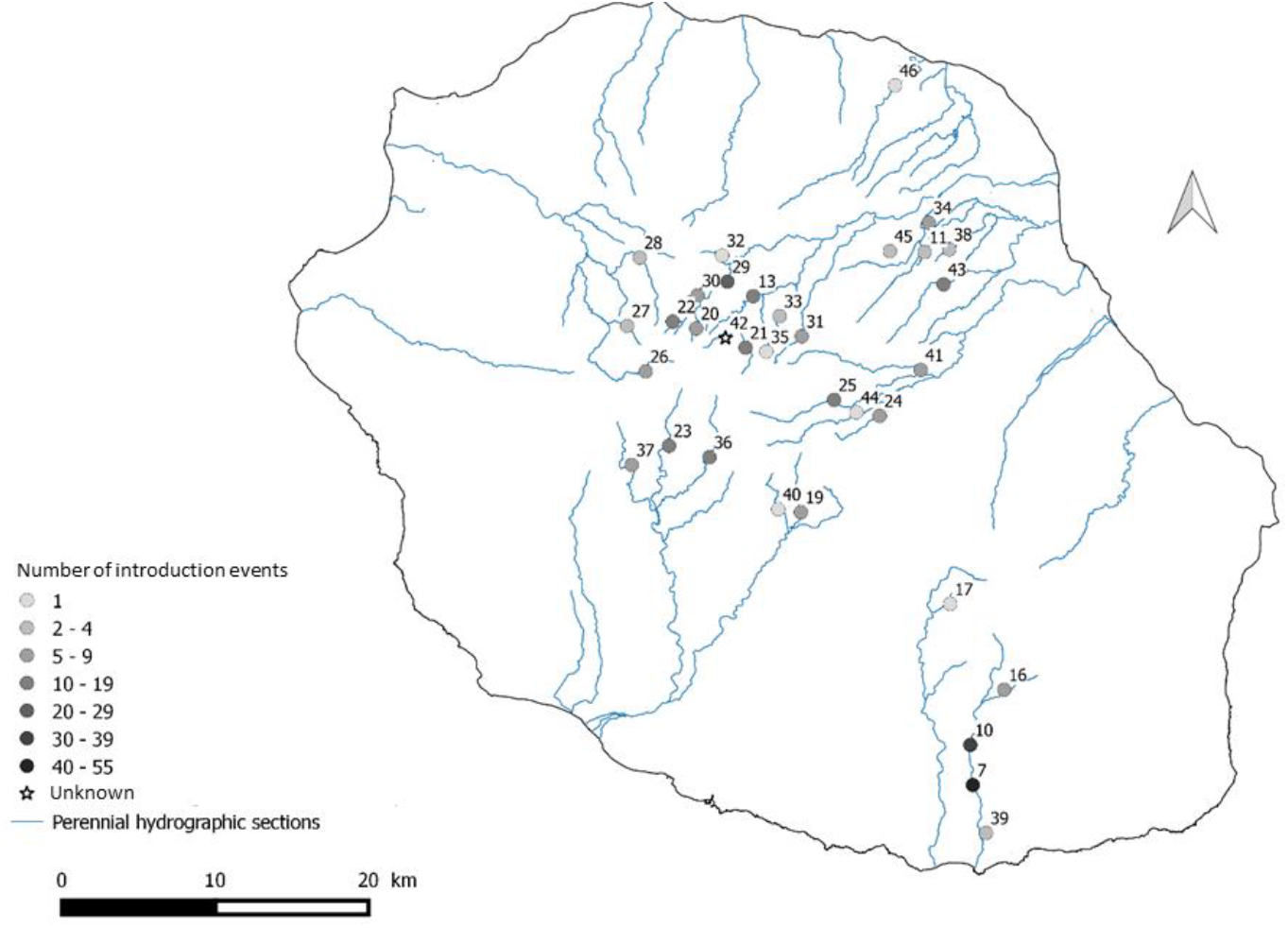
Map of Reunion Island showing perennial rivers and the location and introduction effort at 33 sites where rainbow trout were introduced from 1940-2019. The number of introduction events, used as a proxy of the effort, is represented by increasing shades of gray. The star symbol represents an introduction site with unknown effort. See Table 2 for more information about the sites.

### Seasonal water temperature profiles

Mean monthly water temperatures at the study sites ranged greatly, from 11.0-22.1°C in winter and 14.6-27.7°C in summer (Fig. 3). Seasonality, which can be described by a decrease in temperature in winter, was distinct at some sites, while temperature profiles remained flat throughout the year at several other sites. Self-sustaining populations of trout were present at sites with low water temperature and high seasonal variability, but some sites with low water temperature had no trout, despite trout introduction. At sites where trout were present, mean temperature in summer and winter were 17.5°C (standard deviation (SD) = 0.8) and 14.9°C (SD = 1.0), respectively. The warmest month on record at a site with trout reached a mean temperature of 19.6°C (SD = 1.04) (site 42 in March 2020).

**Fig. 3.**
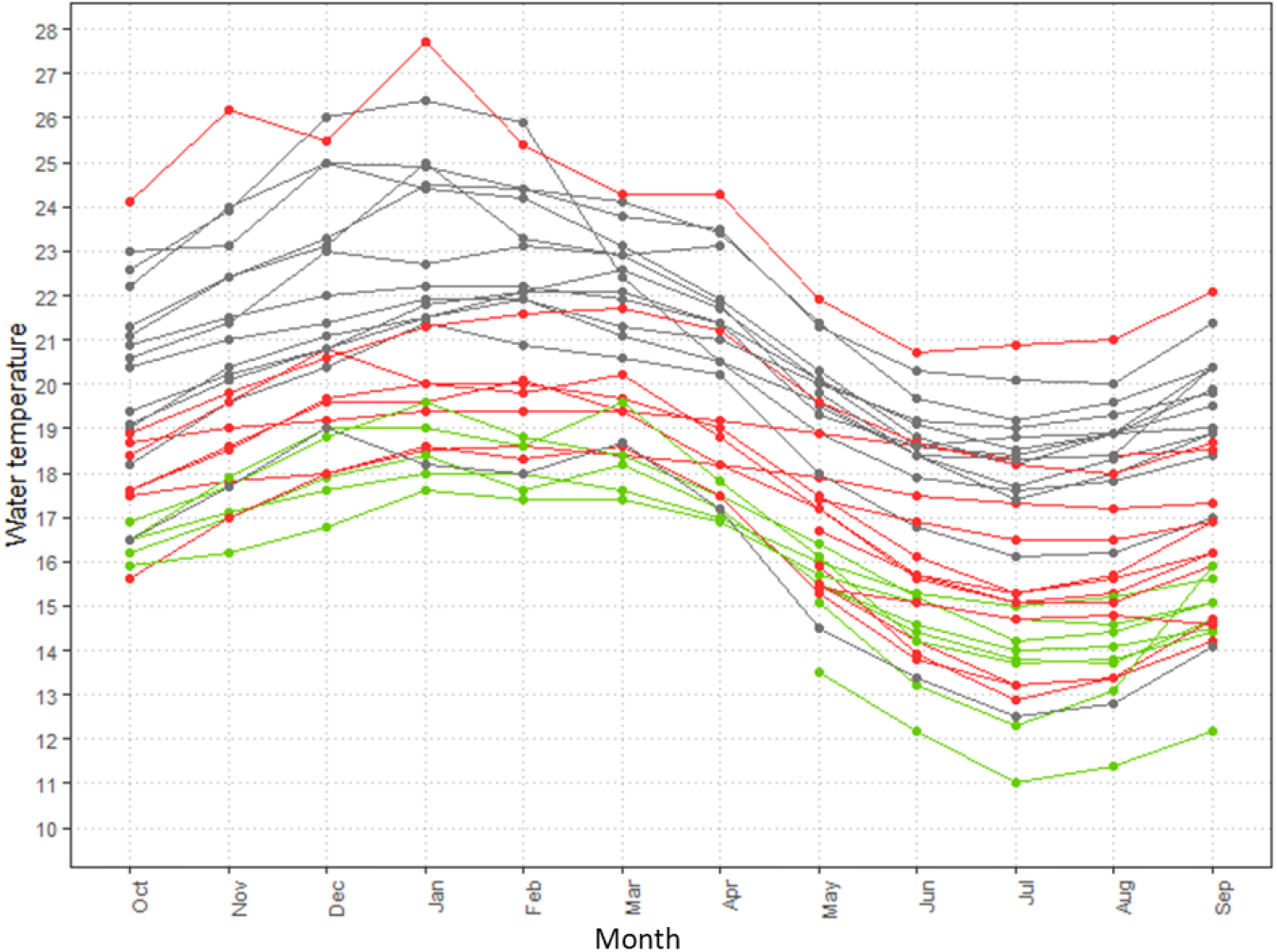
Mean monthly water temperatures at 35 sites on perennial rivers on Reunion Island. Colors show the sites where attempts to introduce rainbow trout resulted in self-sustaining populations (green) or were not successful (red), and sites with no introduction events (gray).

### Using elevation as a proxy of temperature

Mean monthly water temperature correlated strongly with both elevation and season (Table 3), which explained as much as 76.6% of the variance in the data. Adding the interaction between elevation and season did not improve the fit of the model, which indicated that the mean decrease in water temperature for each meter in elevation was similar in winter and summer (Table 3, Fig. 4).

**Table 3.** Modeling mean monthly water temperature on Reunion Island as a function of elevation and season.

| Model | np | AIC | ΔAIC | Pseudo-R<sup>2</sup> |
| --- | --- | --- | --- | --- |
| Temp ~ 1 | 1 | 5775.1 | 4426 | 0.000 |
| Temp ~ Elevation | 2 | 2886.5 | 1537.4 | 0.500 |
| Temp ~ Elevation + Season | 3 | 1349.1 | 0.00 | 0.766 |
| Temp ~ Elevation × Season | 4 | 1349.7 | 0.6 | 0.766 |

**Fig. 4.**
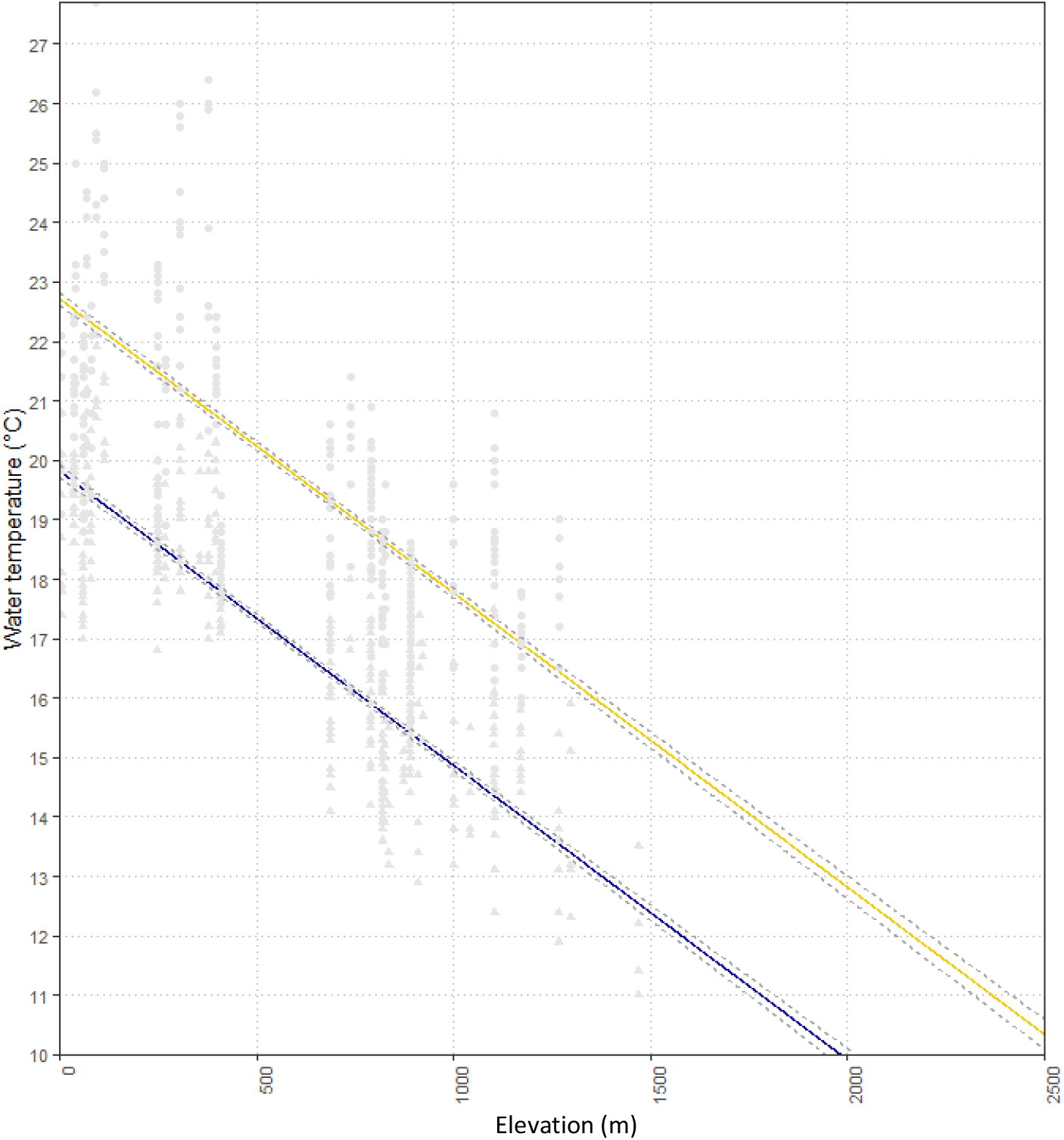
Relationship between mean monthly water temperature and elevation in winter (blue) and summer (yellow) on Reunion Island. The solid lines are model estimates (Table 3), and the gray circles are raw data.

## DISCUSSION

Rainbow trout is a stenotherm species that is native to temperate rivers of North America. It has been extensively introduced all over the world and the literature review indicates that tropical regions are no exception (Fig. 1 and S1). Despite the recognized plasticity of trout, its long-term persistence in the tropics seems challenging due to its affinity for cold water. On Reunion Island, the introduction of trout into at least 35 different sites in river stretches across the country illustrates a long-term and widely spread introduction effort across high-elevation perennial rivers. Despite the recorded release of > 400,000 rainbow trout, we are aware of only 10 self-sustaining populations.

We found no evidence of the presence of self-sustaining populations outside sites where trout had been introduced. If trout disperse, they likely do so over very short distances. Several environmental factors can hinder trout expansion across rivers on Reunion Island. First, the island’s topography, with steep mountains and many natural waterfalls that result in a high level of habitat fragmentation, strongly limits rainbow trout dispersal from introduction sites. Temporary drying of river stretches due to the highly permeable substrate also limits dispersal. We do not know when the populations established, but based on the long history of introduction (> 80 years), it is likely that some populations have sustained themselves for more than one decade. It is also possible that self-sustaining populations went extinct and re-established themselves at different times in recent history, since trout were introduced into rivers regardless of whether or not they were already present. Nevertheless, there is no evidence for a large expansion of trout from introduction sites, and the risk of expansion from current self-sustaining populations seems low.

The observation of only 10 sites with self-sustaining populations, along with the absence of expansion from these sites, suggest that the environmental conditions that trout encounter after introduction or while dispersing may not meet their ecological requirements well. High water temperature may limit the presence of trout in perennial rivers of Reunion Island, since rainbow trout are sensitive to temperature (Fausch 2007). These tropical rivers have a wide range of water temperatures, with higher variability among sites (spatial variability) than within sites (seasonal variability). As expected, the warmest sites had no trout, while the coldest sites did have them. On Reunion Island, trout experience a mean monthly temperature of 17.5°C in summer, with a maximum of 19.6°C.

This range differs from the optimal temperature range for rainbow trout defined in the literature (i.e. 10-16°C (Raleigh 1984; Jalabert & Fostier 2010)) but lies below the extreme of 30°C recorded in some pools in southern California, United States (Sloat & Osterback 2013). Some of the coldest sites, however, had no trout despite many introduction events. In the trout’s native range, increasing summer temperature is a main concern for its persistence under ongoing climate change. It is predicted that higher summer temperatures will decrease the area of thermally suitable riverine trout habitats and cause them to shift upstream (Ruesch *et al*. 2012; Carlson *et al*. 2017; Isaak *et al*. 2018). In contrast, winter temperature requirements are nearly always met in temperate regions (Sloat & Osterback 2013; Brewitt & Danner 2014). Scott and Poynter (1991) suggested that high winter temperatures, rather than high summer temperatures, influenced the northern limit of trout distribution in New Zealand. On Reunion Island, the highly fragmented habitat prevents trout from dispersing upstream, or even migrating seasonally to colder and more suitable habitats.

High summer temperatures may be responsible for the inability of trout to establish themselves at sites where they were introduced, since these temperatures approach the physiological limits of trout. In winter, trout require temperatures no higher than 13°C for several months for successful reproduction (Morrison & Smith 1986; Pankhurst *et al*. 1996; Jalabert & Fostier 2010), which makes winter a critical period of the life cycle. However, prolonged cold temperatures rarely occur on Reunion Island (8 out of 843 mean monthly temperatures <13°C). The relative rate of increase in summer and winter temperatures will be critical in determining the persistence and extinction risk of trout populations. This has implications for improving understanding of responses of rainbow trout to climate change in its native range. In the future, winter temperatures could become a limiting factor for trout reproduction.

Although detailed water temperature data are rare, the presence of trout populations at high elevations in the tropics, where water temperatures are usually cooler, was no surprise. However, other key requirements may limit the presence of trout, such as natural conditions (e.g. extreme flow regime, spawning habitat, food resources, thermal refugee) and human forcing (e.g. pollution, poaching). Furthermore, although we can model explain 77% of the variance of temperature based on elevation, highly contrasting geologic and topographic features can generate large differences between watersheds. For instance, the Cilaos region of Reunion Island has warmer water due to the presence of geothermal groundwater (sites 23, 36 and 37); thus, it may not be suitable for trout, despite its higher elevation. In contrast, upwelling of cool groundwater in the United States can form thermal refuges that may become critical habitat that allows trout to persist at higher temperatures (Ebersole *et al*. 2001; Sloat & Osterback 2013; Brewitt & Danner 2014).

## AKNOWLEDGMENTS

We are grateful to A. Métro, G. Vienne, J. Maillot, D. Vitry, G. Hoarau and all those who were involved in collecting the data used in this study (FDAAPPMA of Reunion Island, its associated AAPPMA and the water Agency of Reunion Island). The study was funded in part by the French National Office for Biodiversity (TAC-Reunion project), which funded the salary of AM and JR.

## DECLARATIONS

The study was funded in part by the French National Office for Biodiversity (TAC-Reunion project), which funded the salary of AM and JR.

